# Microbial Rhodopsins are Increasingly Favored over Chlorophyll in High Nutrient Low Chlorophyll waters

**DOI:** 10.1101/2021.03.30.437613

**Authors:** Babak Hassanzadeh, Blair Thomson, Fenella Deans, Jess Wenley, Scott Lockwood, Kim Currie, Sergio E. Morales, Laura Steindler, Sergio A. Sañudo-Wilhelmy, Federico Baltar, Laura Gómez-Consarnau

**Affiliations:** University of Southern California, Los Angeles, CA, United States; University of Otago, Dunedin, New Zealand; National Institute of Water and Atmospheric Research, Dunedin, New Zealand; University of Haifa, Department of Marine Biology, Leon H. Charney School of Marine Sciences, Haifa, Israel; University of Vienna, Dept. of Functional & Evolutionary Ecology, Vienna, Austria; Centro de Investigación Científica y de Educación Superior de Ensenada, Ensenada, BC, México

## Abstract

Microbial rhodopsins are simple light-harvesting complexes that, unlike chlorophyll photosystems, have no iron requirements for their synthesis and phototrophic functions. Here we report the first environmental concentrations of rhodopsin along the Subtropical Frontal Zone off New Zealand, where Subtropical waters encounter the iron-limited Subantarctic High Nutrient Low Chlorophyll (HNLC) region. Rhodopsin concentrations were highest in HNLC waters where chlorophyll-*a* concentrations were lowest. Furthermore, while the ratio of rhodopsin to chlorophyll-*a* photosystems was on average 20 along the transect, this ratio increased to over 60 in HNLC waters. We further show that microbial rhodopsins are abundant in both picoplankton (0.2-3μm) and in the larger (>3μm) size fractions of the microbial community containing eukaryotic plankton and/or particle-attached prokaryotes. These findings suggest that rhodopsin phototrophy could be critical for microbial plankton to adapt to resource-limiting environments where photosynthesis and possibly cellular respiration are impaired.

**Originality-Significance statement:** High Nutrient Low Chlorophyll (HNLC) regimes cover approximately 30% of the global ocean surface and play a crucial role in the Earth’s carbon cycle. Here we show that microbial rhodopsins are particularly abundant in a HNLC region of the Subantarctic ocean, where chlorophyll abundance is relatively low and photosynthesis and respiration might be impaired due to iron limitation. These data suggest that rhodopsin phototrophy can contribute significantly to the energy budgets of HNLC regions, capturing meaningful amounts of light that cannot be channeled through photosynthesis.

## Introduction

The Subantarctic region of the Southern Ocean is a crucial regulator of the global climate system. Not only is it one of the largest sinks of atmospheric CO2 in the world ocean (Metzl et al., 1999), but also the source of nutrients that ultimately fuel most of the global ocean’s primary production through the convective formation of intermediate waters in this region (Toggweiler et al., 1991; Sarmiento et al., 2004). For that reason, the importance of chlorophyll-based photosynthesis in the Subantarctic waters to the global carbon budget has been studied for decades (Boyd et al., 1999; Doblin et al., 2011). Yet, the magnitude of rhodopsin-based phototrophy and its role in sustaining microbial communities in the Subantarctic HNLC oceanographic regime remains unknown. Microbial rhodopsins are light-driven ion-pumps (Béjà et al., 2000) present in microorganisms of all life domains, with notably over 80% of the surface marine bacteria containing these genes (Dubinsky et al., 2017; Sieradzki et al., 2018).

The simplicity and low synthesis cost of rhodopsin photosystems compared to chlorophyll further suggest that they have a critical role in light energy acquisition, particularly in resource deplete environments where photosynthesis is limited (Raven, 2009). Supporting this hypothesis, several culture studies have shown that rhodopsin phototrophy can improve bacterial growth, survival, or reduce respiration rates when labile organic matter resources are scarce (Gómez-Consarnau et al., 2007; 2010; Steindler et al., 2011). Also, in situ, the abundance of rhodopsin genes appears to be negatively correlated to chlorophyll and inorganic nutrient concentrations (Campbell et al., 2008). To date, ambient rhodopsin concentrations (i.e., the number of rhodopsin photosystems per volume of seawater) have only been reported for the Mediterranean Sea and the Eastern Atlantic Ocean, where they also tended to be inversely related to phytoplankton biomass, nitrate, and phosphate concentrations (Gómez-Consarnau et al., 2019). Despite this empirical evidence, not all studies on rhodopsin phototrophy reflect a clear association with macronutrients or organic matter availability, suggesting that other regulating factors may exist (Pinhassi et al., 2016). For instance, a study in the Chesapeake Bay found that the percent of rhodopsin-containing cells was positively correlated to salinity using a microscopy-based method (Keffer et al., 2015; Maresca et al., 2018). Therefore, expanding rhodopsin observations to additional key marine regions is essential to further elucidate their regulation mechanisms and overall importance globally.

The relationship between rhodopsin phototrophy and iron availability has not been tested thoroughly, and never in the HNLC Subantarctic waters where microbial growth is limited by the availability of this trace element (Sedwick et al., 1999; Sander et al., 2015). Unlike chlorophyll-based photosynthesis, rhodopsin phototrophy does not involve any known redox reactions, and its functioning is independent of electron carriers such as iron (Raven, 2009). Rhodopsin-like genes have been found in populations of the diatom *Pseudo-nitzschia granii* from the North Pacific HNLC region, and further culture studies revealed that the highest levels of both rhodopsin transcripts and proteins occur under low iron conditions in a *P granii* strain (Marchetti et al., 2015). This suggests that this diatom may, indeed, rely on rhodopsin phototrophy when iron concentrations are insufficient to adequately perform photosynthesis. Furthermore, given the high iron requirements of the respiratory electron transport chain, heterotrophic bacterial respiration can also be impaired in HNLC regions due to iron limitation (Tortell et al., 1996).

## Results and Discussion

Here we studied the distributions of the two most important solar energy transducing systems, rhodopsin and chlorophyll-*a* (chl-*a*), on a 60 km transect across the Subtropical Frontal Zone off New Zealand (Munida Microbial Observatory Time Series; MOTS), which traverses through three contrasting oceanographic regimes and water masses: Coastal neritic (CNW), Subtropical (STW), and Subantarctic waters (SAW) (jillett, 1969) (Figure 1A and Supplementary Information). This geographical transect provides a unique opportunity to evaluate the relative contribution of each photosystem in three contrasting environments ranging from coastally-influenced waters to an HNLC iron-limited region (Figure 1B).

**Figure 1.**
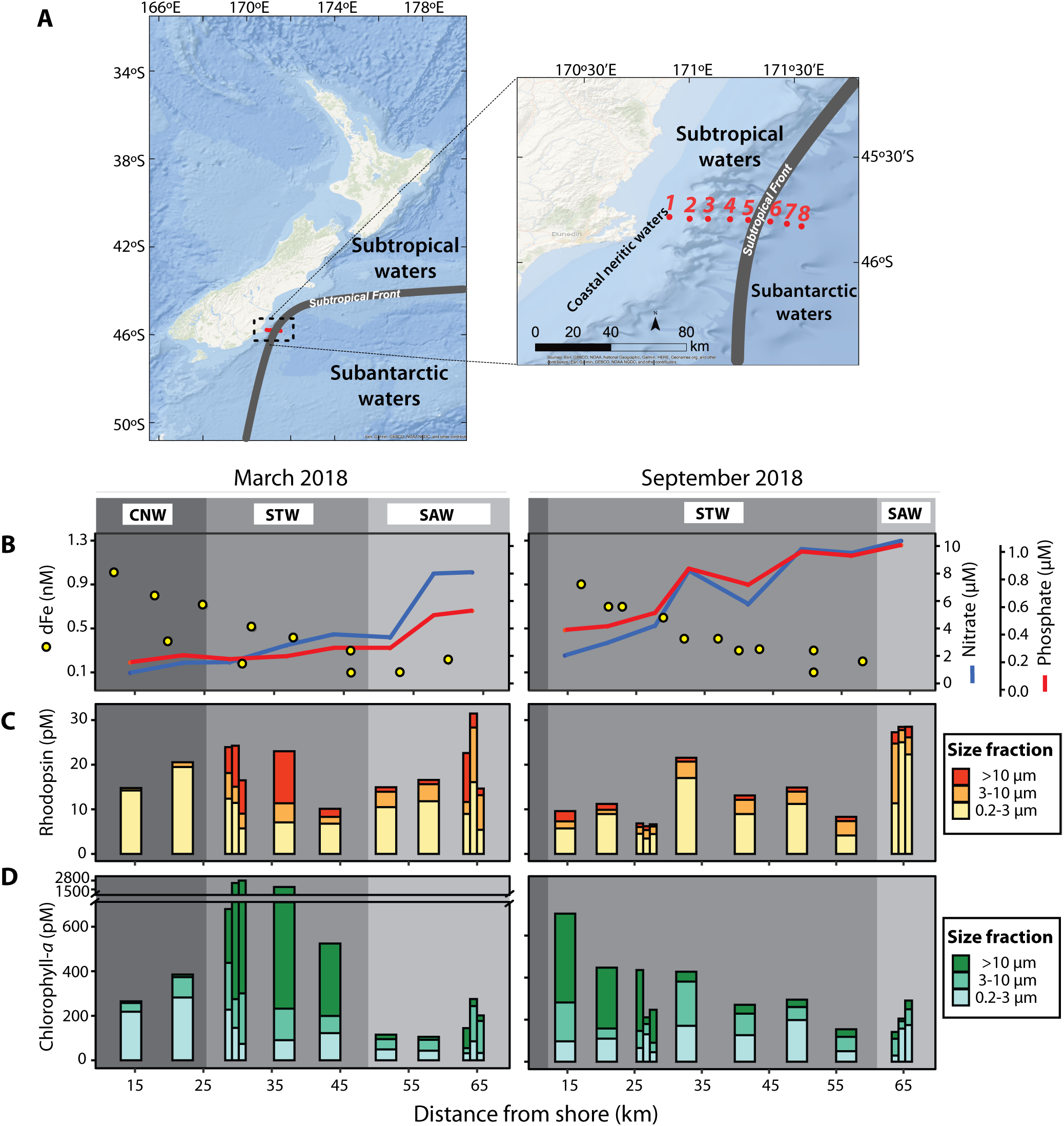
**(A)** Location of the sampling stations (shown in red) along the Munida Microbial Observatory Time Series (MOTS) in Coastal Neritic (CNW), Subtropical (STW); and Subantarctic (SAW) waters. The position of the Subtropical front, which moves seasonally, belongs to March of 2018. Temperature and salinity plots are shown in Figure S1. **(B)** Surface concentrations of dissolved nitrate, phosphate, and iron (dFe) from discrete sampling locations. Nitrate and phosphate concentrations belong to the March and September 2018 cruises. Average dissolved iron concentrations are from previously reported cruises of March and September of years 2000-2003 (Sander et al., 2015). **(C)** Rhodopsin and **(D)** chlorophyll-a levels in picoplankton (0.2-3.0 μm), nanoplankton (3.0-10.0 μm), and >10.0 μm size fractions of the microbial community. The narrower columns in panels C-D represent photosystem concentrations from samples collected at three different depths, 2m, 20m, and deep chlorophyll maximum (from left to right).

Surface rhodopsin and chl-*a* concentrations displayed particular spatial and temporal trends along the transect; rhodopsin ranged from 7 to 27 pM (Figure 1C) while chlorophyll varied substantially more (16-fold; 100-1600 pM; Figure 1D). The highest chl-*a* concentrations were found in different size fractions, depending on the sampling station and season. Rhodopsins were present in all size fractions of the microbial community, with more than 35% of the signal being found in the large size fractions (>3μm). Although rhodopsin genes and transcripts had previously been identified in large microbial communities (>0.8μm) in a temperate estuary (Maresca et al., 2018), these are the first data reporting actual rhodopsin quantifications in nano- or micro-plankton in any open-waters marine system, indicating their presence in eukaryotes and/or particle-attached prokaryotes. However, most of the total rhodopsin content (62%) was found in the picoplankton fractions. These observations, together with rhodopsin gene data from the global ocean (Pinhassi et al., 2016), suggest that rhodopsin phototrophy is widespread in marine microbial communities and that it is primarily a prokaryotic light-capturing feature.

Rhodopsins reached their highest concentrations (~30 pM) at the SAW HNLC stations, coinciding with the lowest chl-*a* levels (100-140 pM; Figures 1CD, 2A), suggesting an increased prevalence of rhodopsin-containing microbial plankton or cellular rhodopsin quotas in the HNLC region. The vertical distributions of rhodopsin (Figure 2A) revealed a maximum above the deep chlorophyll maximum, with less vertical fluctuations than chl-*a*. Within a given depth profile, chl-*a* concentrations fluctuated between 2-4-fold, while rhodopsin levels varied <2-fold. The depth distributions of these photosystems are consistent with previous observations (Gómez-Consarnau et al., 2019) and with the notion that rhodopsin synthesis is energetically advantageous only at high irradiances (Kirchman and Hanson, 2013).

**Figure 2.**
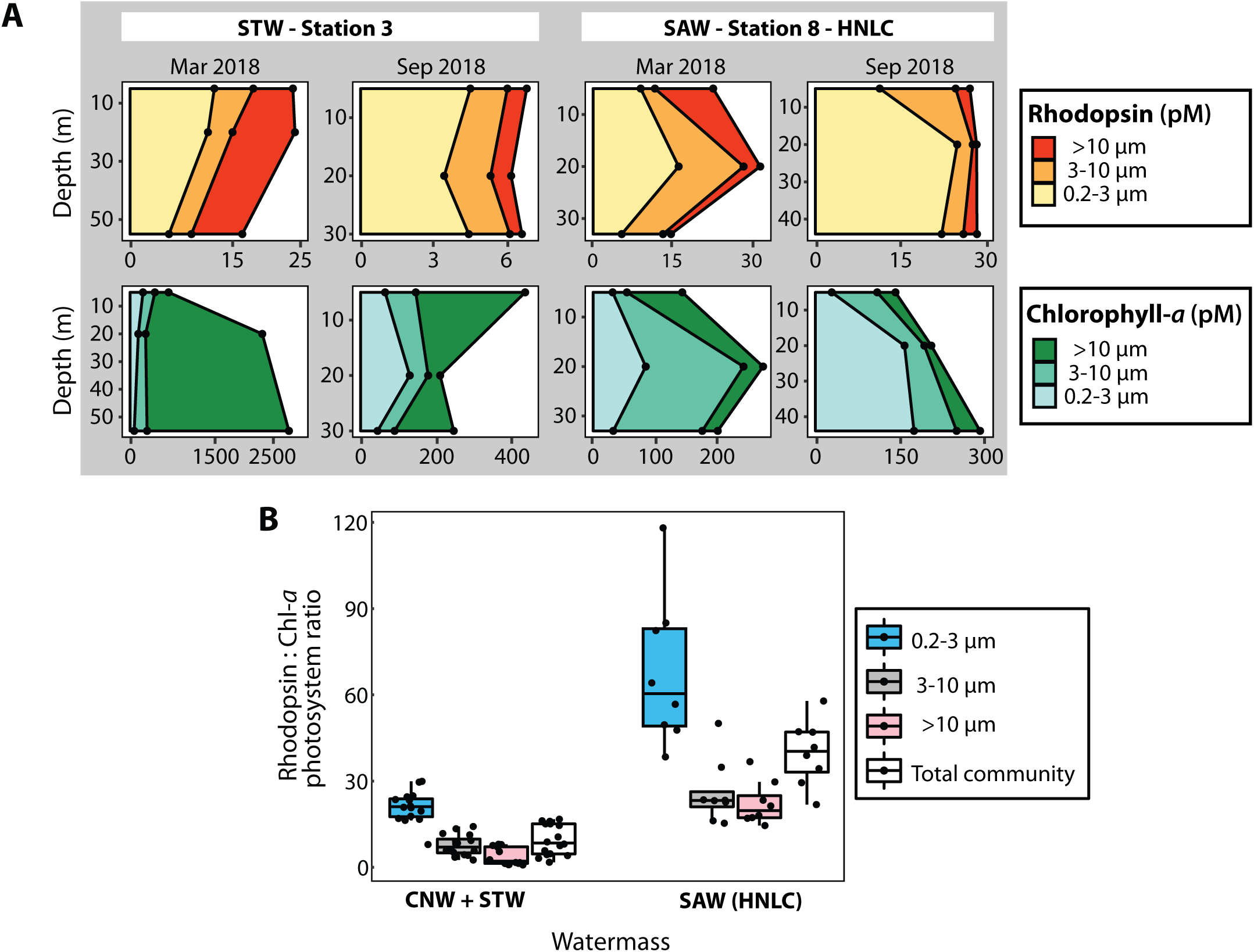
**(A)** Depth profiles of rhodopsin and chlorophyll-a concentrations measured in March and September of 2018. Station-3 and station-8 were located in Subtropical (STW) and Subantarctic (SAW) waters, respectively. **(B)** Rhodopsin to chlorophyll-*a* photosystem ratios in different size fractions of the microbial community, grouped by water mass. Photosystem abundance was calculated assuming one molecule of the retinal for rhodopsin and 300 molecules of chlorophyll-*a* for each chl-*a* photosystem (Mirkovic et al., 2016). Rhodopsin:Chlorophyll-*a* ratios were significantly higher in the HNLC Subantarctic compared to Coastal neritic and Subtropical water masses (Welch two sample t-test, p=6.9·10-4 for 0.2-3 μm, p=1.1·10-3 for 3-10 μm, p=6.0·10-5 for >10 μm size fractions, and p=3.3·10-5 for the entire community).

Rhodopsin photosystem abundances were on average 20 times higher than those of chl-*a* throughout the transect. Yet in HNLC waters, the ratio of rhodopsin to chl-*a* photosystems (R:C ratio) was significantly higher than in the other regions (40±11 compared to 10±5; Figure 2B), suggesting an increase in photoheterotrophy over photosynthesis associated with water mass characteristics (Figure 1B). Similarly, Marchetti et al. (2015) reported an increase in the relative abundance of rhodopsin transcripts and proteins for the diatom *P. granii* under iron deplete conditions. Our observations in natural microbial communities of this HNLC region suggest that the increased R:C photosystem ratios could also be caused by physiological changes within the cells, as observed in P. granii. However, we cannot rule out an increase due to the presence of a larger number of rhodopsin-utilizing organisms.

Picoplankton displayed the highest R:C ratios, increasing 3-fold from 21±5 in the STW and CNW to 68±21 in the SAW-HNLC region (Figure 2B). Thus, while there are several known microbial strategies for coping with iron stress (e.g., the production of siderophores (Tagliabue et al., 2017); high affinity transporters and decreased cell size (Sunda and Hunstman, 1997)), our data show that HNLC environments appear to be selectively enriched with microbial communities that can cope with iron stress by harvesting sunlight through rhodopsins when photosynthesis and/or respiration are compromised. In fact, the metabolic versatility gained through rhodopsin photoheterotrophy may explain the relatively similar bacterial abundances and respiration rates previously reported in STW and SAW despite substantial differences in iron availability (Baltar et al., 2015). Given that respiration is the most iron-demanding process and the primary energy-generating mechanism in heterotrophs (Raven, 1988), the increase in R:C among picoplankton suggests that rhodopsin energy capture provides an ecological advantage during iron limitation. Furthermore, it implies that there is an additional and still unknown amount of solar energy fueling HNLC ecosystems that needs to be considered in energy budgets. Identifying the major rhodopsin-containing microbial groups (both heterotrophs and eukaryotic phytoplankton) in HNLC regions will be central to elucidate the potential ecological processes impacted by this metabolism. Finally, revealing the intricacies of rhodopsin phototrophy as a coping mechanism against resource limitation is likely to reshape our understanding of energy acquisition and the present and future carbon cycle in the ocean.

## Materials and Methods

Seawater samples were collected from 8 stations along the MOTS transect at several depths on March 26 and September 20, 2018 (Figure 1; Supplementary Information). Surface seawater (2m depth) was sampled at all stations, whereas the deep chlorophyll maximum (DCM) was only sampled at stations 3 and 8. Approximately 10-liter samples were size-fractionated by an in-line serial filtration system (10 μm, 3μm, and 0.22 μm pore sizes) using a peristaltic pump. Rhodopsin and chlorophyll-based photosystem concentrations were determined using retinal and chlorophyll-*a* as proxies. Both pigments were extracted from the filters with methanol (Garrido and Roy, 2015; Gómez-Consarnau et al., 2019). Aliquots from each extract were used for immediate fluorometric chlorophyll-*a* quantification using the non-acidification method (Knap et al., 1994) while retinal concentrations were determined through LC/MS/MS (Gómez-Consarnau et al., 2019). Rhodopsin photosystem abundances were then estimated assuming that each rhodopsin photosystem contains one molecule of retinal (Larkum et al., 2018) and that each chlorophyll-based photosystem contains on average 300 molecules of chl-*a* (Mirkovic et al., 2016). Total rhodopsin and chl-*a* concentrations were calculated by adding the three size fractions. The specific LC/MS/MS operational conditions were slightly modified from Gómez-Consarnau et al. (2019) according to Kane and Napoli (2010) to include the use of an internal standard (see Supplementary Information).

Total dissolved iron (dFe) concentrations (operationally defined as the fraction that passed through a 0.45 μm filter) used in this study were obtained from previous March and September cruises and are reported in Sander et al. (2015). The samples for dFe were stored at a pH <2 and quantified after a dithiocarbamate organic extraction (Bruland et al., 1979).

## Acknowledgments

We would like to thank the captain and crew of R/V Polaris II, and Hannah Adams for her assistance with pigment extractions. This research was supported by Grant No OCE1924464 from the United States National Science Foundation (NSF) to L.G.-C. and by Grant No 2019612 from the United States-Israel Binational Science Foundation (BSF) to L.S., as well as the Rutherford Discovery Fellowship to F.B.

## Author contributions

B.H., F.B., and L.G-C. designed research; B.H., B.T., F.D., J.W., S.L., K.C., S.E.M., L.S., S.A.S-W, F.B. and L.G.-C. performed research and analyzed data. B.H. and L.G.-C. wrote the manuscript with the input of all authors.

